# Improving the Quality of Co-evolution Intermolecular Contact Prediction with DisVis

**DOI:** 10.1101/2022.11.03.515006

**Authors:** Siri C. van Keulen, Alexandre M.J.J. Bonvin

## Abstract

The steep rise in available protein sequences and structures has paved the way for bioinformatics approaches to predict residue-residue interactions in protein complexes. Multiple sequence alignments are commonly used in intermolecular contact predictions to identify co-evolving residues. These contacts, however, often include false positives (FPs), which may impair their use to predict three dimensional structures of biomolecular complexes and affect the accuracy of the generated models.

Previously, we have developed DisVis to identify false positive data in mass spectrometry cross-linking data. DisVis allows to assess the accessible interaction space between two proteins consistent with a set of distance restraints. Here, we investigate if a similar approach could be applied to co-evolution predicted contacts in order to improve their precision prior to using them for modelling complexes.

In this work we analyze co-evolution contact predictions with DisVis in order to identify putative FPs for a set of 26 protein-protein complexes. Next, the DisVis-reranked and the original co-evolution contacts are used to model the complexes with our integrative docking software HADDOCK using different filtering scenarios. Our results show that HADDOCK is robust with respect to the precision of the predicted contacts due to the 50% random contact removal during docking and using DisVis filtering for low precision contact data. DisVis can thus have a beneficial effect on low quality data, but overall HADDOCK can accommodate FP restraints without negatively impacting the quality of the resulting models. Other more precision-sensitive docking protocols might, however, benefit from the increased precision of the predicted contacts after DisVis filtering.

## Introduction

What is the prediction quality of protein complexes for which isolated structures are available but protein-protein interface (PPI) information is not? Unfortunately, there is still a low probability of predicting (or identifying) the correct PPI in those cases and this has been one of the main challenges for the structural bioinformatics field for the past decades. The steady increase of protein sequences in data banks such as Uniprot^1^ and major technical advances in the structural biology field^2^ have been important factors for the enhanced prediction accuracy of protein complexes over the past year^3^. With the rise in experimental data, software is now being developed to leverage the large quantity of sequences and structures by mining them, via co-evolution or machine learning (ML) algorithms^4–6^, for example. The release of Alphafold2^6^ has demonstrated that ML approaches can compete or even outperform the state-of-the-art software packages in the protein structure-prediction field^7^. Besides protein structures, recent studies are also exploring Alphafold2’s predictive power for protein-protein^8,9^ and protein-peptide^10,11^ complexes.

Co-evolution has been proven to be an important tool to identify residues at potential PPIs^12^. Identifying co-evolving residue pairs requires the availability of multiple sequence alignments (MSA) of orthologous sequences. When applied to the prediction of intermolecular contacts, an additional complexity comes from predicting the correct pairing of the sequences of the two proteins when considering multiple paralogues. The predicted intermolecular contacts derived from co-evolving residues can then be used in *de-novo* modeling of protein complexes^5,13^. Although this technique has mainly been used for prokaryotic systems, recent findings suggest eukaryotic complexes could also benefit from applying co-evolution prediction approaches^12,14,15^.

Independent of the protein system, one major challenge in co-evolution predictions remains the presence of false positive (FP) contacts. Although FP contacts are deduced from MSAs in the same way as correct contacts, they do not describe the physiological protein-protein interface. When such contact data are used to model protein complexes, these false positives can negatively affect the modeling results as they potentially steer the model away from the correct solution, reducing the prediction accuracy. This is a more general problem, which, for example, also occurs in cross-linking mass-spectrometry (XL-MS) data which also suffer from FPs. To deal with this problem we have previously developed DisVis, available both as a web server^16^ and python package ^17^, which, given the 3D structures of the component of a complex, can assess the accessible interaction space defined by a set of distance restraints and identify possible false positives. Similarly, the identification and removal of FPs in co-evolution predicted contacts through DisVis could potentially improve the modelling of protein complexes based on residue-residue contact information.

Here, we use 26 protein-protein complexes and co-evolution contact predictions selected from the work of Green et al.^18^ to evaluate whether DisVis analysis can help in FP removal. We then assess the impact of using this information (original co-evolution or DisVis-filtered contacts) on the quality of the docking results using our integrative modelling software HADDOCK which allows defining distance restraints to guide the docking. We show that DisVis-filtering increases the precision of the predicted contacts and that HADDOCK is not very sensitive to this precision increase in the predicted contacts as it is able to generate correct models even in the presence of a significant number of FP contacts.

## Methods

### Dataset Preparation

The study by Green et al.^18^ was used to extract 26 protein dimers together with their respective top 20 co-evolution predicted interface contacts obtained through EVcomplex^5^ (Fig. 1). The protein complexes were selected according to the total number of true residue-residue contacts predicted within the top 10 intermolecular co-evolution contacts of each system, information extracted from the supporting material of Green et al.^18^. We selected cases having a top 10 contact precision ranging from 20 to 100%, ensuring an equal distribution over contact precision ranges (Table 1, Supporting Fig. 1): Five complexes per total number of true contacts (2, 4, 6, 8 or 10) were included. An additional complex with 9 true contacts was added, resulting in a total number of 26 complexes. Besides the distribution in true contacts, the resolution of the X-ray structures was taken into consideration by including the highest resolution structures possible and avoiding redundancy. The structures of the monomers were prepared for use in DisVis and HADDOCK using a python script to rename the protein chains (chain A and B) with pdb-tools^19,20^ (`pdb_chain and pdb_tidy`). While those structures have exactly the same backbone conformation as in the experimental reference complex, their side chains were perturbed and optimized using SCWRL4 by Green et al. ^18^, which thus represent a semi-unbound conformation for docking purposes.

**Figure 1.**
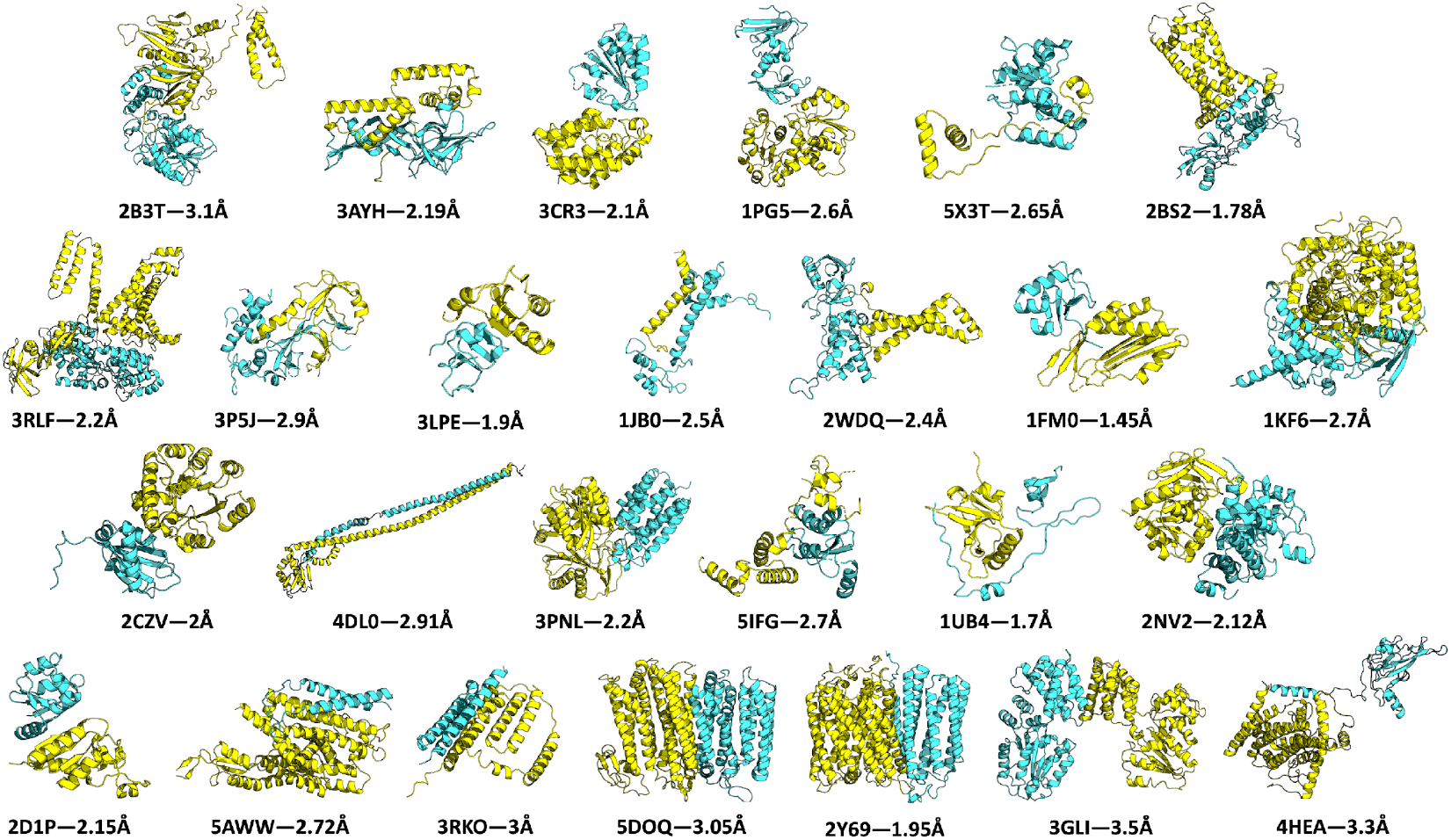
Dataset of 26 dimers used in this study. In each dimer the two chains are highlighted in yellow and blue. The PDB ID as well as the resolution of the experimental structure in Ångstrom are depicted. The representation of the shown protein complexes was obtained by using PyMOL^21^.

**Table 1.** Structure information of the 26 hetorodimeric protein-protein complex dataset used in this study. Each entry describes one dimer with its corresponding PBD ID, the chains that have been used in DisVis and HADDOCK, the resolution of the experimental structure, the Green ID^18^ equivalent, the number of true residue-residue contacts and corresponding top-10 and top-20 precision (%) according to calculations performed by Green et al.^18^, and the number of predicted contacts (*Present Green* 10/20) in the Green top-10 and top-20 for which the corresponding residues are present in the experimental protein structures.

| PDB ID | Chain 1 | Chain 2 | Resolution (Å) | Green ID<sup>a</sup> | Green top 10<sup>b</sup> | Green top 20<sup>b</sup> | Present Green contacts 10/20 |
| --- | --- | --- | --- | --- | --- | --- | --- |
| 3CR3 | B | D | 2.1 | allpdb0993 | 2 (20%) | 2 (10%) | 9/18 |
| 3AYH | A | B | 2.19 | allpdb0972 | 2 (20%) | 3 (15%) | 10/19 |
| 1PG5 | A | B | 2.6 | allpdb0696 | 2 (20%) | 2 (10%) | 10/20 |
| 5X3T | C | D | 2.65 | allpdb1682 | 2 (20%) | 2 (10%) | 8/18 |
| 2B3T | B | A | 3.1 | allpdb1938 | 2 (20%) | 4 (20%) | 9/18 |
| 2BS2 | C | B | 1.78 | allpdb0461 | 4 (40%) | 5 (25%) | 10/20 |
| 3LPE | A | B | 1.9 | allpdb1080 | 4 (40%) | 4 (20%) | 8/16 |
| 3RLF | F | E | 2.2 | allpdb0211 | 4 (40%) | 4 (20%) | 8/18 |
| 1KF6 | M | N | 2.7 | allpdb0234 | 4 (40%) | 4 (20%) | 10/20 |
| 3P5J | C | B | 2.9 | allpdb0336 | 4 (40%) | 4 (20%) | 8/16 |
| 1FM0 | E | D | 1.45 | allpdb0609 | 6 (60%) | 8 (40%) | 10/20 |
| 2CZV | A | C | 2 | allpdb0803 | 6 (60%) | 8 (40%) | 10/20 |
| 2WDQ | C | B | 2.4 | allpdb0144 | 6 (60%) | 9 (45%) | 10/20 |
| 1JB0 | J | F | 2.5 | allpdb0058 | 6 (60%) | 8 (40%) | 10/19 |
| 4DL0 | E | G | 2.91 | allpdb0367 | 6 (60%) | 10 (50%) | 9/19 |
| 1UB4 | A | C | 1.7 | allpdb0732 | 8 (80%) | 9 (45%) | 10/20 |
| 2NV2 | L | K | 2.12 | allpdb0854 | 8 (80%) | 12 (60%) | 10/20 |
| 2D1P | D | F | 2.15 | allpdb0190 | 8 (80%) | 11 (55%) | 10/20 |
| 3PNL | A | B | 2.2 | allpdb1128 | 8 (80%) | 9 (45%) | 10/20 |
| 5IFG | B | A | 2.7 | allpdb1601 | 8 (80%) | 11 (55%) | 10/19 |
| 5AWW | Y | G | 2.72 | allpdb0550 | 9 (90%) | 18 (90%) | 10/20 |
| 2Y69 | N | P | 1.95 | allpdb0089 | 10 (100%) | 13 (65%) | 10/20 |
| 3RKO | F | E | 3 | allpdb0153 | 10 (100%) | 15 (65%) | 10/19 |
| 5DOQ | A | B | 3.05 | allpdb2088 | 10 (100%) | 18 (90%) | 10/20 |
| 4HEA | Q | O | 3.3 | allpdb1728 | 10 (100%) | 15 (65%) | 10/20 |
| 3GLI | F | J | 3.5 | allpdb1822 | 10 (100%) | 11 (55%) | 10/20 |

a) The unique ID number of each complex used by Green et al.<sup>18</sup> in their supplementary information.
b) Number of true contacts in the top 10 and top 20 co-evolution contacts according to the definition used by Green et al.<sup>18</sup>, using the experimental structures and a heavy-atom interface cutoff of 8 Å calculated with haddock-tools (https://github.com/haddock/haddock-tools) to identify the true contacts. The contact precision is shown in brackets.

### DisVis Scoring of Co-evolution Predicted Intermolecular Contacts

For each dimer within our dataset, the top-20 co-evolution predicted contacts (see Data availability) were used as input for DisVis through its web server implementation (https://wenmr.science.uu.nl/disvis). The two monomer structures together with a list of predicted intermolecular contacts were submitted with the *complete scanning option* settings (1Å voxel size and 9.72° scanning angle). Predicted residue-residue contacts that involved residues absent in the available 3D protein structures were removed from the co-evolution contact lists prior to DisVis calculations (see Contact Precision and Table 1: *Present contacts*). The upper distance limit for the co-evolution contacts was set to 10 Å between Cα atoms (in their work, Green et al. used 8 Å between Cβ-Cβ atoms) as during the rotational scan only Cα atoms are considered, which was implemented to reduce computational costs^17^. The DisVis calculated z-scores were used to rank the residue-residue contacts.

The z-score is calculated for each distance restraint by taking into account each DisVis modelled complex which meets at least one of the distance criteria included by the user. For each complex that meets this requirement, all violated restraints are calculated and stored. This results in a violation matrix in which the violation data of all approved complexes are combined. Each row of the matrix represents the number of consistent restraints, from 1 to N, and each column describes the frequency of restraint violation per distance restraint in which at least N restraints are consistent. This violation matrix is used to calculate the z-score per restraint:

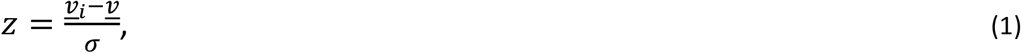

where 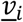 is the average per column *i* of the violation matrix, and 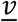 and *σ* describe the violation matrix average and standard deviation, respectively^17^. The resulting z-scores were ordered for this study from low (negative z-score) to high (positive z-score), least to most likely to be a false positive.

From the DisVis-reranked co-evolution contacts, the top 10 and 5 were extracted for use as distance restraints in HADDOCK. The entire set of 20 contacts (i.e. without DisVis filtering) was also considered as well as DisVis filtered data, using a z-score threshold of 0.5 or 1.0.

### Docking Protocols

The docking calculations were performed using a local installation of HADDOCK 2.4. The docking protocol in HADDOCK consists of three stages^22^. In the first stage (it0), rigid body docking is performed (it0) with the distance restraints defined between the two chains guiding the docking. From the 1000 (default) generated models, the top 200 based on the it0 HADDOCK score progress to the next step. The second stage (it1) consists of a semi-flexible simulated annealing in torsion-angle space during which flexibility at the interface is introduced step wise, first along the side chains and later for both side chains and backbone. By default, the flexible interface is defined automatically for each model from an analysis of residues that are in close contact between the chains. All structures from it1 are transferred to the final step of the docking protocol (itw) which consists in HADDOCK 2.4 of a final energy minimization (previous versions of HADDOCK were performing a very short optimization by molecular dynamics simulation in explicit solvent – this option is still available but turned off by default in version 2.4). Finally, the models are scored based on the HADDOCK itw scoring function which is a linear combination of energetic terms:

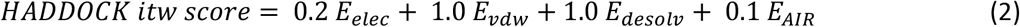

where E_*elec*_ and E_*vdw*_ correspond to the electrostatics and Van-der-Waals intermolecular energies, respectively, E_*desolv*_ to the desolvation energy and E_*AIR*_ represents the energy term assigned to the Ambiguous Interaction Restraints (AIRs) (in this case the predicted contacts)^22^.

Default settings were used for all HADDOCK runs^22^, except for the random removal of restraints (see Table 2). The DisVis-reranked distance restraints or the original 20 co-evolution contacts were included as input in the ambiguous restraints class. Nine different docking protocols (Table 2) were performed with HADDOCK, which differ in the number and type of restraints considered and the percentage of restraints randomly discarded for each model. The latter option makes HADDOCK potentially less sensitive to wrong (e.g. false positive) restraints. Intermolecular co-evolution distance restraints were defined as distances of 3 (lower bound) to 7 Å (upper bound) between Cβ atoms of the two chains, except for glycine residues for which the Cα atom was selected. This definition of the distance restraints is the same as in the docking calculations performed by Green et al.^18^. Besides co-evolution restraints, additional intramolecular Cα-Cα distance restraints were included during docking for protein chains in which parts of the protein structure were missing to keep the domains together during the refinement stage (note that this is done automatically in the web server). These restraints were calculated with the `restrain_bodies.py` script from haddock_tools (https://github.com/haddocking/haddock-tools).

**Table 2.** The different HADDOCK protocols tested with the original or DisVis-reranked co-evolution restraints.

| Protocol | Number of Restraints | Prediction Method | Random Removal<sup>a</sup> (%) | Selection based on DisVis z-score | ID |
| --- | --- | --- | --- | --- | --- |
| 1 | 20 | Co-evolution<sup>b</sup> | 50 | - | EV20-50 |
| 2 | 10 | Co-evolution<sup>b</sup> | 0 | - | EV10-0 |
| 3 | 10 | DisVis reranked<sup>c</sup> | 0 | - | DisVis10-0 |
| 4 | 10 | Co-evolution<sup>b</sup> | 50 | - | EV10-50 |
| 5 | 10 | DisVis reranked<sup>c</sup> | 50 | - | DisVis10-50 |
| 6 | 5 | Co-evolution<sup>b</sup> | 0 | - | EV5-0 |
| 7 | 5 | DisVis reranked<sup>c</sup> | 0 | - | DisVis5-0 |
| 8 | $\leq 20^d$ | DisVis filtered | 50 | $< 0.5^d$ | DisVis20-50<0.5 |
| 9 | $\leq 20^d$ | DisVis filtered | 50 | $< 1.0^d$ | DisVis20-50<1 |

a) Percentage of random removal of restraints. This random removal is done for each model calculated.
b) Co-evolution intermolecular contacts directly taken from Green et al.<sup>18</sup>.
c) Co-evolution intermolecular contacts taken from the DisVis reranking.
d) Co-evolution intermolecular contacts taken from the DisVis reranking by applying a z-score cutoff: z-score lower than 0.5 or 1.0, for protocol 8 and 9, respectively. The number of contacts selected varies for the protein complexes between 8 and 17 for the 0.5 cutoff, and 12 and 20 for the 1.0 cutoff.

### Analysis

#### Contact Precision

The contact precision was calculated for each complex as a function of the number of contacts considered (based on the original or DisVis rankings). The precision 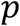 was defined as the number of true contacts divided by the total number of contacts considered:

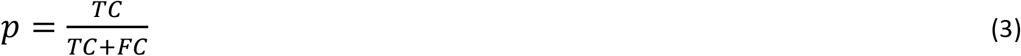

where TC stands for true contact, a contact of which both residues are present at the interface of the reference complex within an 8 Å distance cutoff of each other, considering all heavy atoms. False contacts (FC) are those for which the shortest distance between any heavy atoms exceeds this 8Å cutoff. Subsequently, the average contact precision 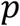 was calculated over all complexes.

#### Interface Root-Mean Square Deviation and Success Rate Calculation

The quality of each complex was determined by calculating the interface root-mean-square deviation (i-RMSD), which is obtained by aligning the backbone atoms at the protein-protein interface of both protein chains on the reference complex, using all residues making contacts within a 10 Å cutoff with the partner molecule. The quality of each model is rated according to the Critical Assessment of Predicted Interactions (CAPRI) with an i-RMSD of ≤ 1 Å denoted as high, ≤ 2 Å as medium and ≤ 4 Å as acceptable quality^23^. We did not consider the fraction of native contacts in this study since in our experience with HADDOCK the limiting factor for defining the quality of a model is the i-RMSD (i.e. a model will never be “downgraded” in quality because of a lower fraction of native contact value).

These model quality ratings are used to calculate the success rates per tested condition. The success rate is defined as the percentage of targets for which a model of acceptable (or better) quality has been generated within the top N (N=1, 5, 10, 20, 50, 100 and 200) ranked models based on the HADDOCK itw score.

## Results

The 26 protein complexes (Fig. 1) used in this study are taken from the dataset published by Green et al.^18^ with 2, 4, 6, 8 and 10 true contacts in the top-10 co-evolution predicted contacts according to the true contact definition used by Green. Five complexes for each number of true contacts were included in our dataset as well as one additional complex with 9 true contacts (Table 1). The number of true contacts in the top-20 for these 26 complexes ranges from 2 to 18 (precision of 10% to 90%) (Table 1). The co-evolution intermolecular contacts from Green et al.^18^ were reranked by DisVis via their z-score to identify potential false positives from the predicted contacts. Different selections of co-evolution restraints were tested (Table 2) to assess the impact of contact precision on the docking performance and model quality.

### Reranking Predicted Co-evolution Contacts with DisVis Enhances the Precision of the Top 10

Co-evolution intermolecular contacts produced by EVcomplex form the starting point for the DisVis analysis of this study. Twenty co-evolution contacts per complex were assessed by DisVis and reranked according to their obtained z-score (see Methods). Subsequently, the DisVis-reranked contacts were compared to the original co-evolution results. The average contact precision (Fig. 2) shows that a difference in precision is already present between the co-evolution and DisVis-reranked contacts in the top 1 (precision of DisVis-reranked 88±32% versus 81±40% for the original contacts). The precision of the co-evolution contacts also decreases faster from the top 1 to the top 10 than for the DisVis-reranked contacts. For the top 10 contacts, the difference in precision is 6% (DisVis-reranked 67%±29% versus 61%±27% for the original contacts). Including more contacts, up to the maximum of 20 considered, lowers the precision further to 47%.

**Figure 2.**
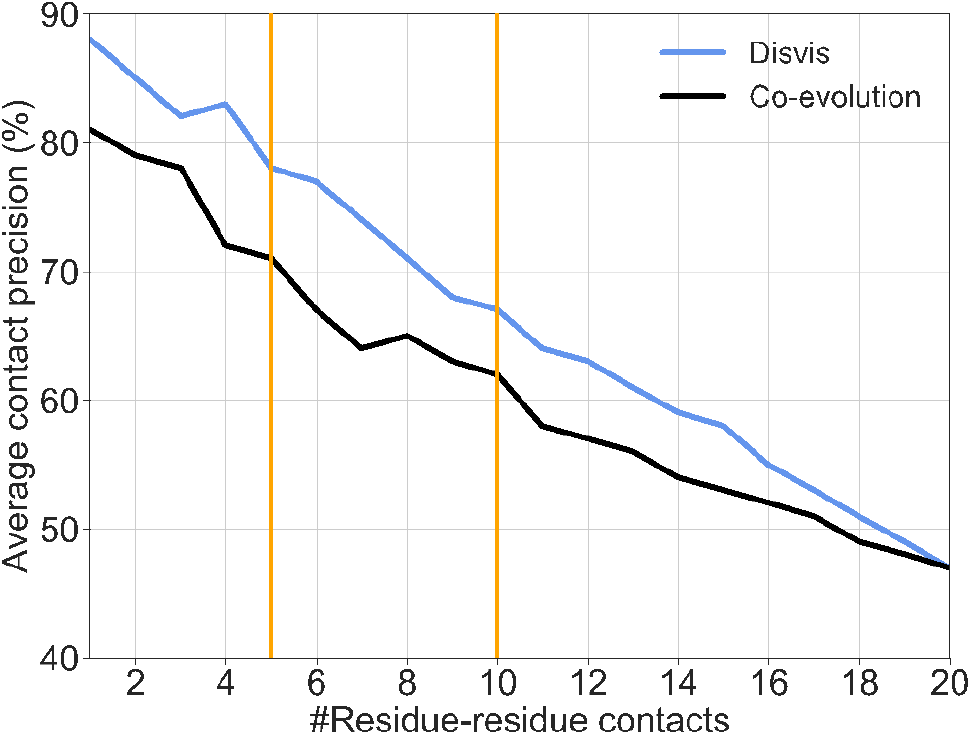
Residue-residue contact precision. Average precision of co-evolution and DisVis-reranked residue-residue contacts calculated over a dataset of 26 dimers. The average precision of the co-evolution predicted contacts are represented by a black line while the DisVis result is shown in blue. Orange lines highlight the two top cutoffs used as input for docking calculations.

### The Number of Contacts Considered rather than their Precision Enhances HADDOCK’s Performance

In order to test the impact of DisVis reranking on the quality of the models generated by HADDOCK, two contact list cutoffs were used as input for docking calculations: top 5 and top 10 (indicated by orange vertical lines in Fig. 2), both using the original EVcomplex ranking and the DisVis reranking of contacts. In addition, as a reference, docking was performed using the original top-20 EVcomplex predictions. The success rates for these different sets of restraints (Table 2) are shown in Figure 3, calculated over the 200 HADDOCK-ranked models after final refinement (itw) (see Methods). Four docking conditions were tested: using the top 5 contacts (EVcomplex or DisVis-reranked) as distance restraints without random contact removal (i), the top 10 contacts (EVcomplex or DisVis-reranked) without random removal of contacts (ii), the top 10 contacts (EVcomplex or DisVis-reranked) with a 50% random removal of provided contacts (iii), and as a reference the top 20 contacts (EVcomplex) with 50% random contact removal (iv) (see Methods and Fig. 3). The random removal of restraints is done per model (1000 models are generated per docking run), meaning that models will be generated based on different combinations of restraints within a docking run.

**Figure 3.**
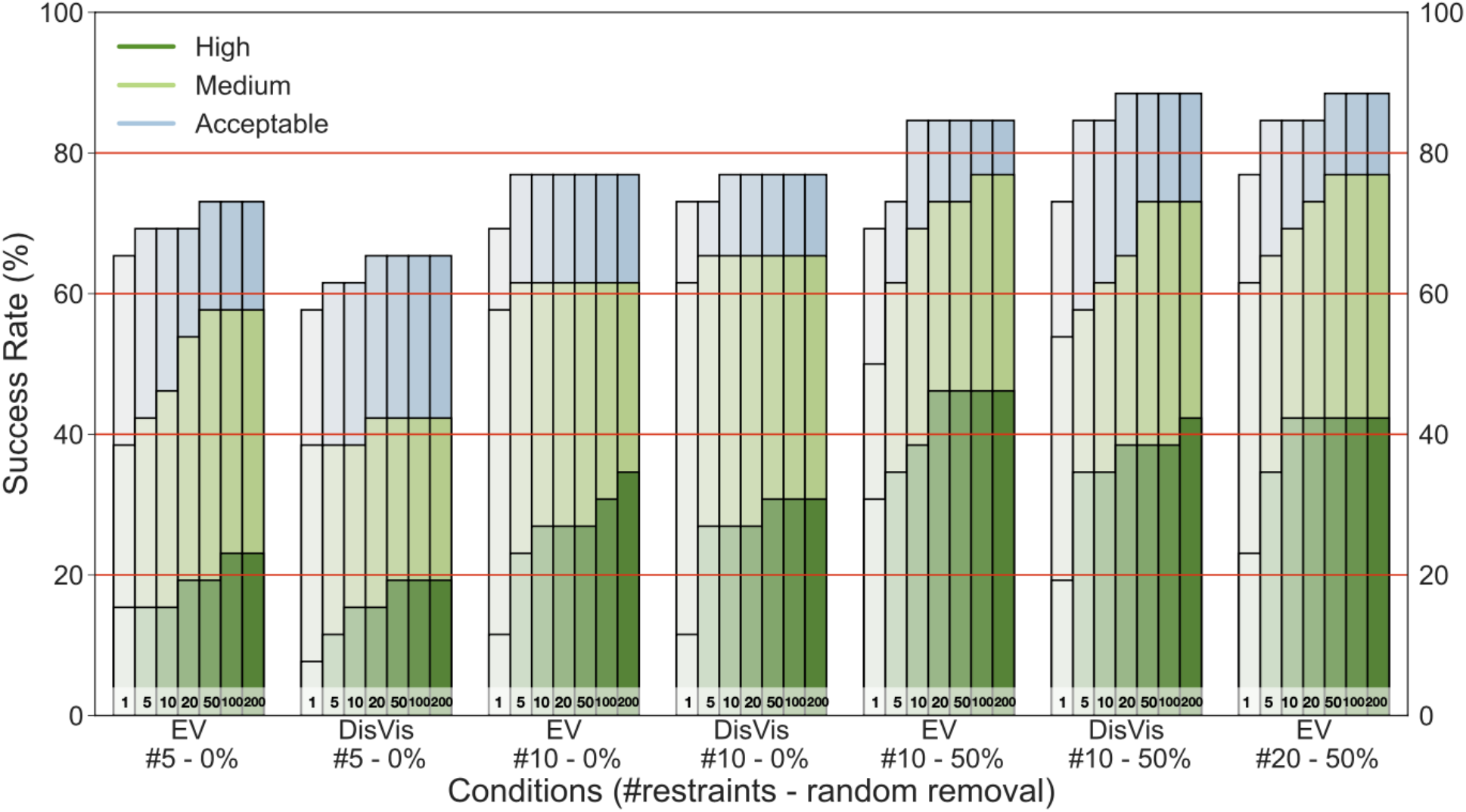
Comparison of co-evolution and DisVis-reranked docking success rates for the 26 dimers dataset. Success rate of co-evolution and reranked DisVis contact lists used as input for protein-protein docking. Three sets of contact lists, 5, 10 and 20, were used to assign distance restraints in HADDOCK. When using the top 5 contacts, all 5 contacts were included in the docking protocol. Hence no random removal was applied. For the top 10, 50% of the included contacts were randomly removed upon docking in #10 - 50% and none were removed in #10 - 0%. The fourth condition represents the docking results using 20 distance restraints with 50 % random removal. Seven bars have been plotted per condition, denoting the top1, 5, 10, 20, 50, 100 and 200 structures according to the HADDOCK itw score. The assignment of a high, medium or acceptable label to a protein complex represents its accuracy in iRMSD with high being ≤ 1 Å (dark green), medium ≤ 2 Å (light green) and acceptable ≤ 4 Å (light blue) (Supplementary Table 1).

The first condition with 5 restraints and no random restraint removal (EV5-0 and DisVis5-0) includes a set of contacts with the highest contact precision compared to the top-10 and top-20 contacts. EV5-0 and DisVis5-0 perform similarly well in the top-10 success rate for high- and medium-quality models. However, EV5-0’s predictions surpass the DisVis setup when it comes to the percentage of acceptable quality predictions. Even though the accuracy of the top-5 restraints used in these protocols is significantly higher than the top-10 contacts for both EVcomplex and DisVis-reranked setups (Fig. 2), EV5-0 and DisVis5-0 are outperformed by the other protocols.

Next, the top-10 contacts were included in four protocols (DisVis10-0, DisVis10-50, EV10-0 and EV10-50) to investigate the impact of random removal of restraints on the docking performance. The DisVis10-50 and EV10-50 protocols (10 restraints and 50% random removal) achieve the best performance with respect to the setups without random removal, reaching an acceptable or higher quality success rate of 85% for the top-10 HADDOCK-scored models. A similar trend can be observed when a cluster-based analysis is performed (Supporting Fig. 2). Hence, turning on random restraint removal (the default setting) improves the docking performance (Fig. 3), making HADDOCK robust to the presence of false positives. The DisVis-reranked performance was also compared to the EVcomplex results, both with random removal of restraints turned on (DisVis10-50 vs EV10-50). When considering the top-5 predicted models, the success rate of the high- and medium-quality models between the two setups is comparable. However, DisVis outperforms the EVcomplex restraints with a success rate of 85% versus 73% for the number of acceptable models in the top 5, suggesting DisVis-reranking can have a quality enhancing effect on co-evolution predicted data when used for docking.

However, none of the top-5 and top-10 DisVis-reranked or EVcomplex setups outperform the EVcomplex condition using 20 restraints and 50% random removal (Fig. 3). The inclusion of 20 restraints during docking results in 35% high-quality structures, 65% medium and 85% acceptable models according to the CAPRI criteria. This finding suggests that although an accuracy improvement within the top-5 and top-10 residue-residue contacts due to filtering with DisVis improves the input data for HADDOCK, using a lower precision contact list with more contacts actually outperforms shorter contact lists with higher precision (Fig. 4 and Supporting Fig. 4). Therefore, HADDOCK appears to be robust with respect to contact precision and benefits from contact quantity.

**Figure 4.**
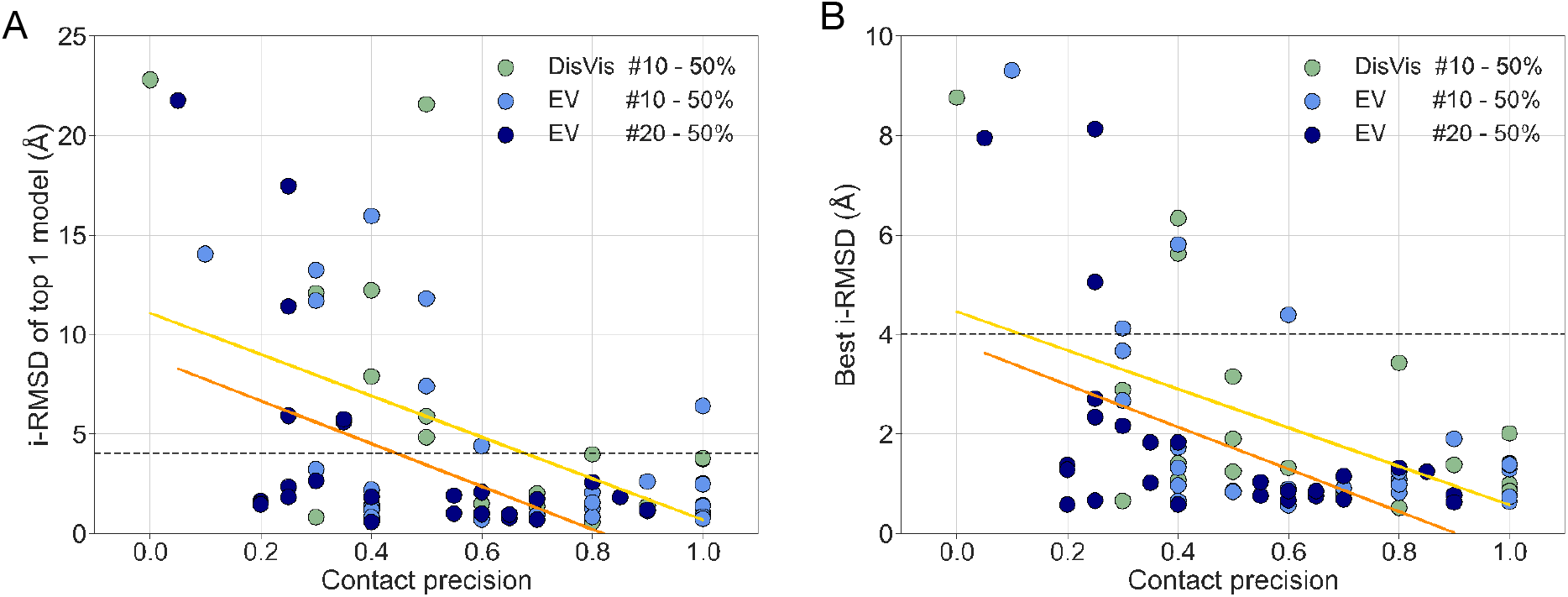
Contact precision versus interface root-mean square deviation (i-RMSD). **(A)** Residue-residue contact precision versus the i-RMSD of the top 1 predicted model per complex, using the HADDOCK itw scoring function. The dark blue circles represent the docking results obtained by using the top 20 EVcomplex contact restraints (Pearson correlation of −0.51). The light blue and green data points show the HADDOCK results from the docking runs performed with the top-10 EVcomplex contacts and the top-10 DisVis-reranked contacts with 50% random removal which have a Pearson correlation of −0.51 and −0.56, respectively. The linear regression fit for the light blue data is highlighted in yellow while the orange line describes the fit for the top-20 EVcomplex results. The dashed black line depicts the 4 Ångstrom CAPRI cutoff for docked models with acceptable quality. **(B)** Residue-residue contact precision versus the model with the best i-RMSD per complex, using the HADDOCK itw scoring function. The dark blue circles represent the docking results obtained by using the top-20 EVcomplex contact restraints (Pearson correlation of −0.51). The light blue and green data points show the HADDOCK results from the docking runs performed with the top-10 EVcomplex contacts and the top-10 DisVis-reranked contacts with 50% random removal which have a Pearson correlation of −0.53 and −0.56, respectively. The linear regression fit for the light blue data is highlighted in yellow while the orange line describes the fit for the top-20 EVcomplex results. The dashed black line depicts the 4 Ångstrom CAPRI cutoff for docked models with acceptable quality.

### Better precision does lead to both better quality and better ranking of models

When analyzing the impact of the precision of residue-residue contacts on the quality of the resulting models in terms of i-RMSD values, it becomes apparent that they are correlated. In Figure 4, the docking results of the best performing protocol, EV20-50 (EVcomplex top 20 contact restraints with 50% removal), is shown in dark blue. The performance of the two runners-up protocols, EV10-50 and DisVis10-50, are depicted in light blue/green. We observe a moderate anti-correlation between contact precision and the i-RMSD of the top-1 docked model or the model with the best i-RMSD (correlation coefficients between −0.51 to −0.56 depending on the data set). More interesting is the fact that irrespective of the dataset, we observe that HADDOCK is able to reliably predict acceptable models in the top ranked models, starting around a contact precision of 0.4 (although acceptable models are already obtained in some cases for precisions as low as 0.2) (Fig. 4B). A comparison of Figures 4A and 4B also shows that the ranking of models improves with the precision, with the top models being of acceptable or better quality when the precision reaches 0.5-0.6.

A comparison of the results obtained by using 10 (both DisVis and EV contacts) or 20 contacts (Fig. 4) shows that for a similar precision, having more contacts does lead to better quality models in general (the dark blue points are in most cases lower than the others). This effect is more apparent at lower precisions and can also be observed in the clustered HADDOCK results (Supporting Fig. 3). These findings indicate that HADDOCK is robust with respect to the precision of contacts and benefits from longer (up to 20 here) contact lists, being able to generate and reliably identify acceptable models down to about 30% precision.

## Discussion

In this study, we have investigated the effect of residue-residue contact filtering on protein-protein docking by comparing the docking results of DisVis-reranked contact restraints to the original co-evolution contacts. Because of the available true contact distribution of the studied dataset, we could analyse how contact quality impacts the docking results. These subsets were defined by using the true contact precision in the Green top 10 (see Methods). Of the 26 complexes, ten fall into the low-quality category with 20-40% true contacts in the top 10, and eleven into the high quality category with true contact precision of 80-100%. Unsurprisingly, success rate analysis for these two groups (20-40% vs 80-100%), show that the original co-evolution contacts with 50% random removal performs best for the 80-100% precision category (Supporting Fig. 5). Overall, the random contact removal (enabled by default in HADDOCK) appears to be crucial to counterbalance the presence of false positives as each of the 1000 docking attempts generates a different set of 50% of the contacts, leading to a robust performance of HADDOCK in regard to contact precision (Fig. 3).

The performance enhancing combination of a large set of distance restraints with medium precision and 50% random removal is also shown in Figure 4B. In this graph, the results clearly demonstrate that while overall interface precision is reduced in the dataset for the 20 contacts setup, HADDOCK generates higher quality models with 20 contacts than 10 contacts at the same interface precision. The difference in ranking performance (Fig. 4A) also shows that while 10 contacts appear to require a contact precision of ~60% to predict an acceptable model at the top 1, 20 contacts achieve a similar quality starting from ~40% precision.

We have also investigated if selecting contacts based on a z-score criterion rather than a predefined number of contacts would improve their quality. Removing contacts with a z-score higher than 0.5 results in an average of 10.6±2.7 contacts per complex with an average precision of 68%±29%. Compared to the original top-20 co-evolution set (Supporting Table 2), including an average 19.2±1.2 contacts with an average precision of 48%±25%, this is an improvement in precision of 20% (68%-48%). The same analysis performed on the low- and high-quality subsets separately leads to 8.8±1.0 contacts per complex for the low-quality set with a precision of 51% and 12.6±2.9 for the high-quality set with an average precision of 83% (Supporting Fig. 6 and Supporting Table 2). Compared to the original top-20 co-evolution contacts subsets (Supporting Table 1), this is an improvement in precision of 24% (51%-27%) and 18% (83%-65%) for the low- and high-quality datasets, respectively. Hence, z-score filtering can positively impact the precision of the contact dataset, especially when the contact set has a low precision initially. This is confirmed by the docking results for the low-quality contacts set (20-40%) (Table 1) when only the contacts with a DisVis z-score lower than 0.5 out of the 20 contacts are included (Supporting Fig. 5). While a z-score cutoff of 0.5 improves the average precision of the remaining contacts, a removal of z-score values higher than 1.0 does not seem to be able to filter the contact data sufficiently (Supporting Table 2), resulting in a similar docking performance as the original co-evolution contact set (Supporting Fig. 5).

However, in a real-world scenario of experimental data or co-evolution data for which a complex structure is not available, the quality of the contacts cannot be assessed before docking. Therefore, comparing and/or combining the top-10 HADDOCK itw-scored structures for both identified approaches, using the original contact data (with 50% random removal) and the DisVis-filtered contacts with z-score<0.5 (and 50% random removal), could provide a way to check the consistency of the solutions between the runs and possibly refine all solutions together, discarding the restraints (in this way the score would only reflect the quality of the interface).

## Conclusion

Intermolecular contacts derived from co-evolution analysis provide a valuable source of information to guide the modelling of protein-protein complexes by docking. These can be used to guide the docking process (as done e.g. in HADDOCK) or as filters to score the generated models. The presence of false positives within the predicted contact data can, however, hamper the docking performance, both in terms of quality and number of acceptable models generated. Here, we have shown that DisVis can reduce the number of FPs in co-evolution contact data by taking into consideration the spatial restrictions imposed by protein structures and the defined contacts. This precision enhancement can have a positive effect on the docking results depending on the software and approach used. Although HADDOCK is robust to the presence of false positive contacts and overall benefits most from a large set of interface contacts and 50% random removal of restraints (the default setup) rather than high interface precision for a small set of contacts, other software or approaches might well benefit from improved precision contact data resulting from DisVis filtering, especially if those contacts are used for scoring purposes rather than to guide the docking. While this work concentrated on co-evolution data, the acquired insights should also be relevant for other types of distance-based information.

## Data availability

The dataset used for this research including the raw data and analysis scripts are available at https://github.com/haddocking/contact-filtering (DOI: 10.5281/zenodo.7260708) and an archive containing in addition all the docking models (top 200 refined HADDOCK models), is available at https://doi.org/10.5281/zenodo.7260736.

## Acknowledgements

This work has been done with the financial support of the Dutch Foundation for Scientific Research (NWO) (PPS Technology Area grant 741.018.201)) and the European Union Horizon 2020 project BioExcel (823830).

## Supporting information

**Supporting Figure 1.**
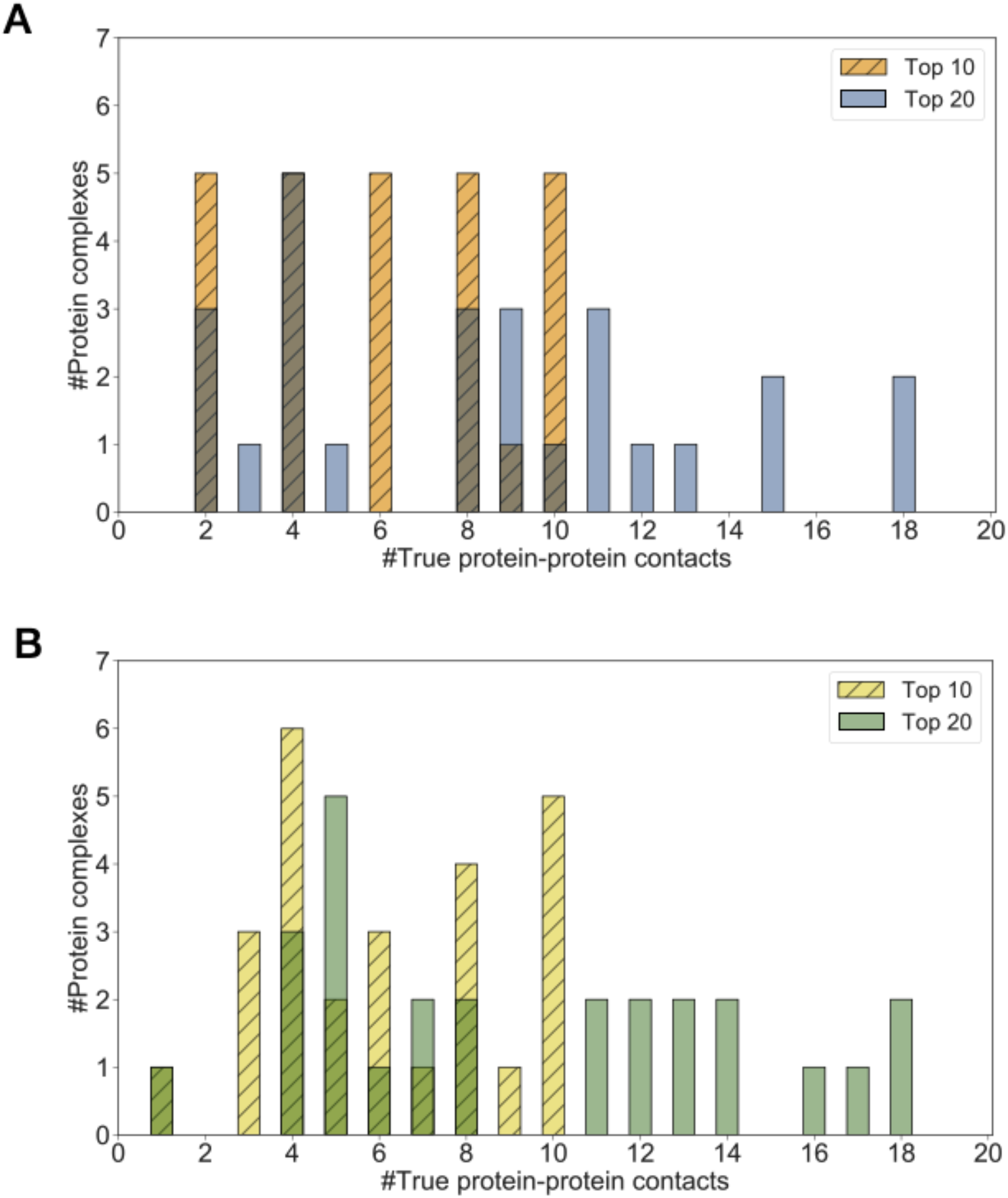
Distribution of true protein-protein contacts of the top 10 and top 20 EVcomplex predictions. A) Contact distribution according to the calculations performed by Green et al.^18^, using an 8 Ångstrom cutoff. B) Contact distribution according to the calculations performed with haddock-tools^24^, using an 8 Ångstrom cutoff.

**Supporting Figure 2.**
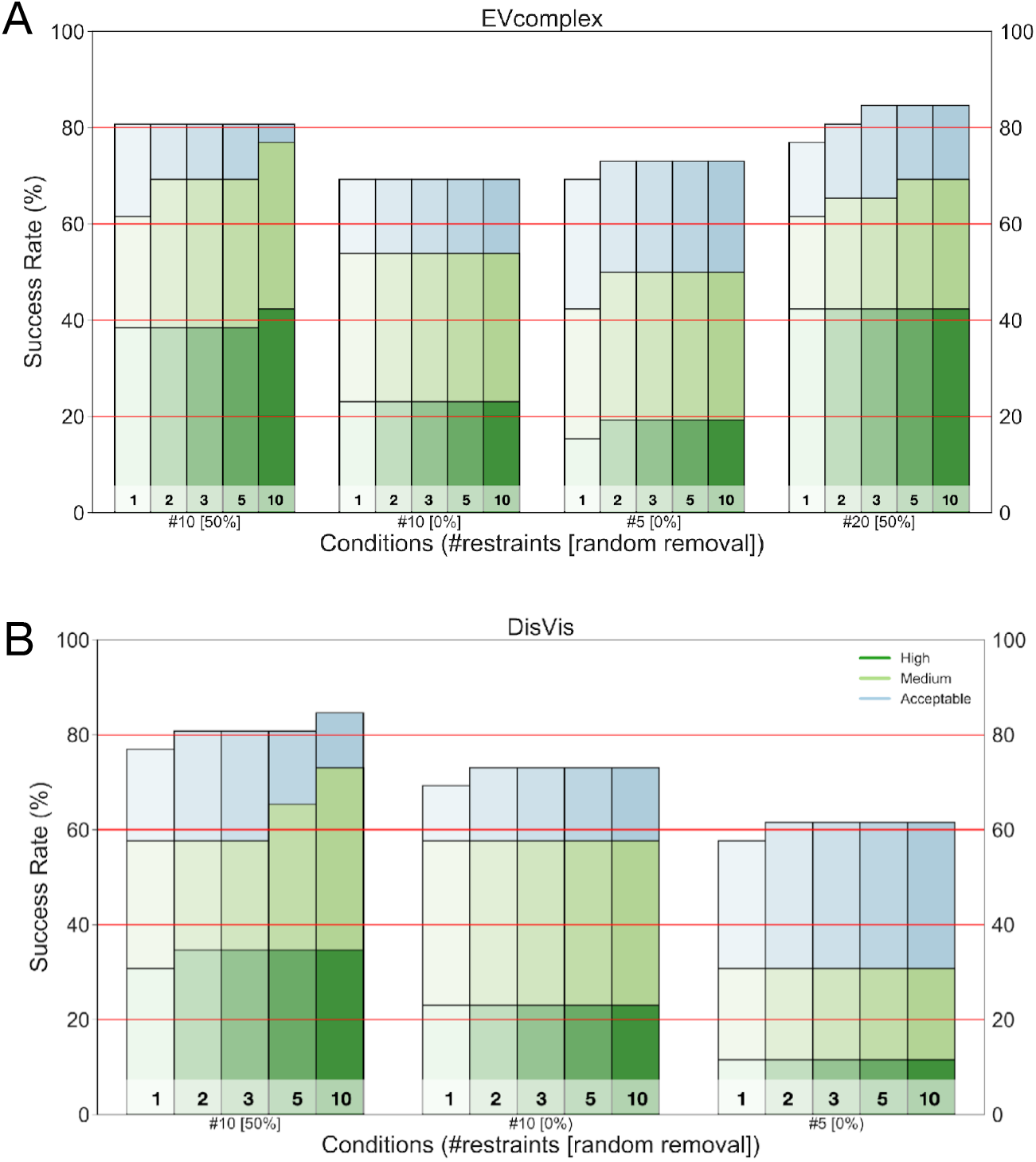
Comparison of co-evolution and DisVis-reranked docking success rates of HADDOCK-scored predicted model clusters. **(A)** Success rate of protein-protein docking using co-evolution contact lists as distance restraints. Three sets of contact lists, 5, 10 and 20, were used to assign distance restraints in HADDOCK. When using the top 5 contacts of the co-evolution prediction, all 5 contacts were included in the docking protocol. Hence no random removal was applied. For the top 10, 50% of the included contacts were randomly removed upon docking in one condition (#10[50%]) and none were removed in #10 [0%]. The fourth condition represents the docking results using 20 distance restraints with 50% random removal. Five bars have been plotted per condition, denoting the top 1, 2, 3, 5 and 10 predicted model clusters according to the HADDOCK itw score. The top four structures in each cluster were included for success rate determination of the 26 dimers. The assignment of a high, medium or acceptable label to a protein complex represents its accuracy in i-RMSD with high being ≤ 1 Å (dark green), medium ≤ 2 Å (light green) and acceptable ≤ 4 Å (light blue). **(B)** Success rate of protein-protein docking using reranked DisVis contact lists as distance restraints. Two sets of contact lists, 5 and 10, were used to assign distance restraints in HADDOCK. When using the top 5 contacts of the DisVis reranked prediction, all 5 contacts were maintained in the docking protocol. Hence no random removal was applied. For the top 10, 50% of the included contacts were randomly removed upon docking in one condition (#10[50%]) and none were removed in #10[0%]. Five bars have been plotted per condition, denoting the top 1, 2, 3, 5 and 10 predicted model clusters according to the HADDOCK itw score. The top four structures in each cluster were included for success rate determination of the 26 dimers. The assignment of a high, medium or acceptable label to a protein complex represents its accuracy in i-RMSD with high being ≤ 1 Å (dark green), medium ≤ 2 Å (light green) and acceptable ≤ 4 Å (light blue).

**Supporting Figure 3.**
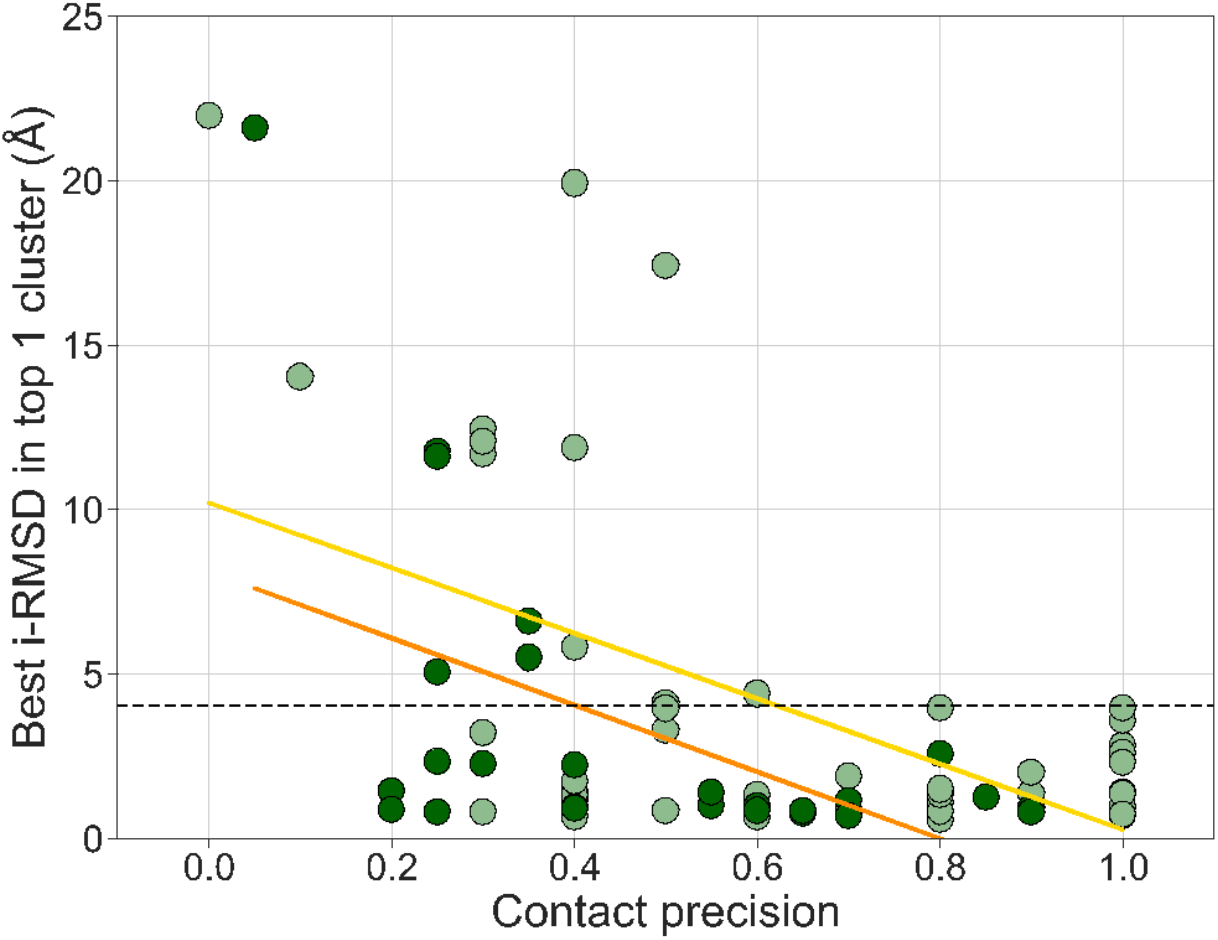
Residue-residue contact precision versus cluster i-RMSD of docking results. Residue-residue contact precision versus the lowest i-RMSD of the top 1 predicted cluster per complex. In order to determine the lowest i-RMSD, the best 4 structures according to the HADDOCK itw score were taken into account. The dark green circles represent the docking results obtained by using the top 20 EVcomplex contact restraints. The light green data show the HADDOCK results from the docking runs performed with the top 10 EVcomplex contacts and the top 10 DisVis-reranked contacts with 50% random removal. The linear regression fit for the light green data is highlighted in yellow while the orange line describes the fit for the top 20 EVcomplex results. The dashed black line depicts the 4 Ångstrom cutoff for the acceptable quality models according to the CAPRI criteria.

**Supporting Table 1.** Average number of acceptable models in it0 (out of 1000), it1 (out of 200) and itw (out of 200) with standard deviations shown in brackets.

| Protocol | # it0 | # it1 | # itw |
| --- | --- | --- | --- |
| EV20-50 | 296.5 (320.3) | 92.3 (72.7) | 92.2 (72.7) |
| EV10-0 | 468.2 (444.3) | 121.8 (91.1) | 121.9 (91.0) |
| DisVis10-0 | 467.0 (443.5) | 122.2 (90.8) | 122.2 (90.7) |
| EV10-50 | 307.2 (310.2) | 87.4 (69.5) | 87.2 (69.5) |
| DisVis10-50 | 307.6 (312.9) | 88.4 (71.2) | 88.5 (71.2) |
| EV5-0 | 451.5 (440.6) | 116.3 (88.3) | 116.5 (88.1) |
| DisVis5-0 | 340.1 (445.4) | 78.0 (88.3) | 77.6 (88.0) |
| DisVis20-50<0.5 | 354.4 (373.4) | 96.6 (79.6) | 96.5 (79.6) |
| DisVis20-50<1 | 346.0 (364.3) | 93.6 (78.4) | 93.6 (78.3) |

**Supporting Figure 4.**
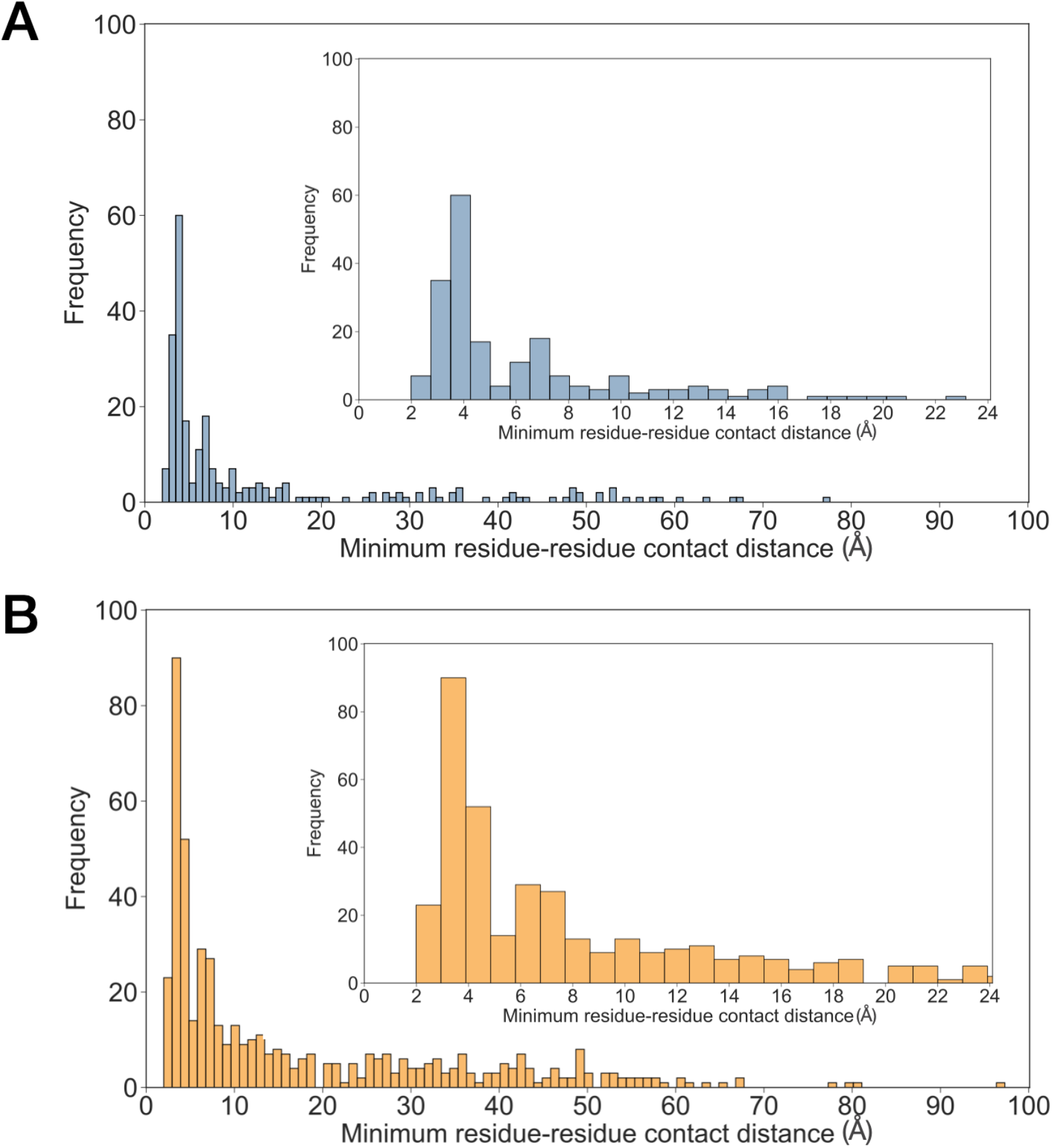
Residue-residue contact distance distribution. Histogram of the EVcomplex top-10 **(A)** and top-20 **(B)** residue-residue contacts for all 26 complexes of the dataset, including their shortest distance. The bin size was set to 100. A zoom of the residue-residue distance distribution from 0-24 Å is depicted as well.

**Supporting Figure 5.**
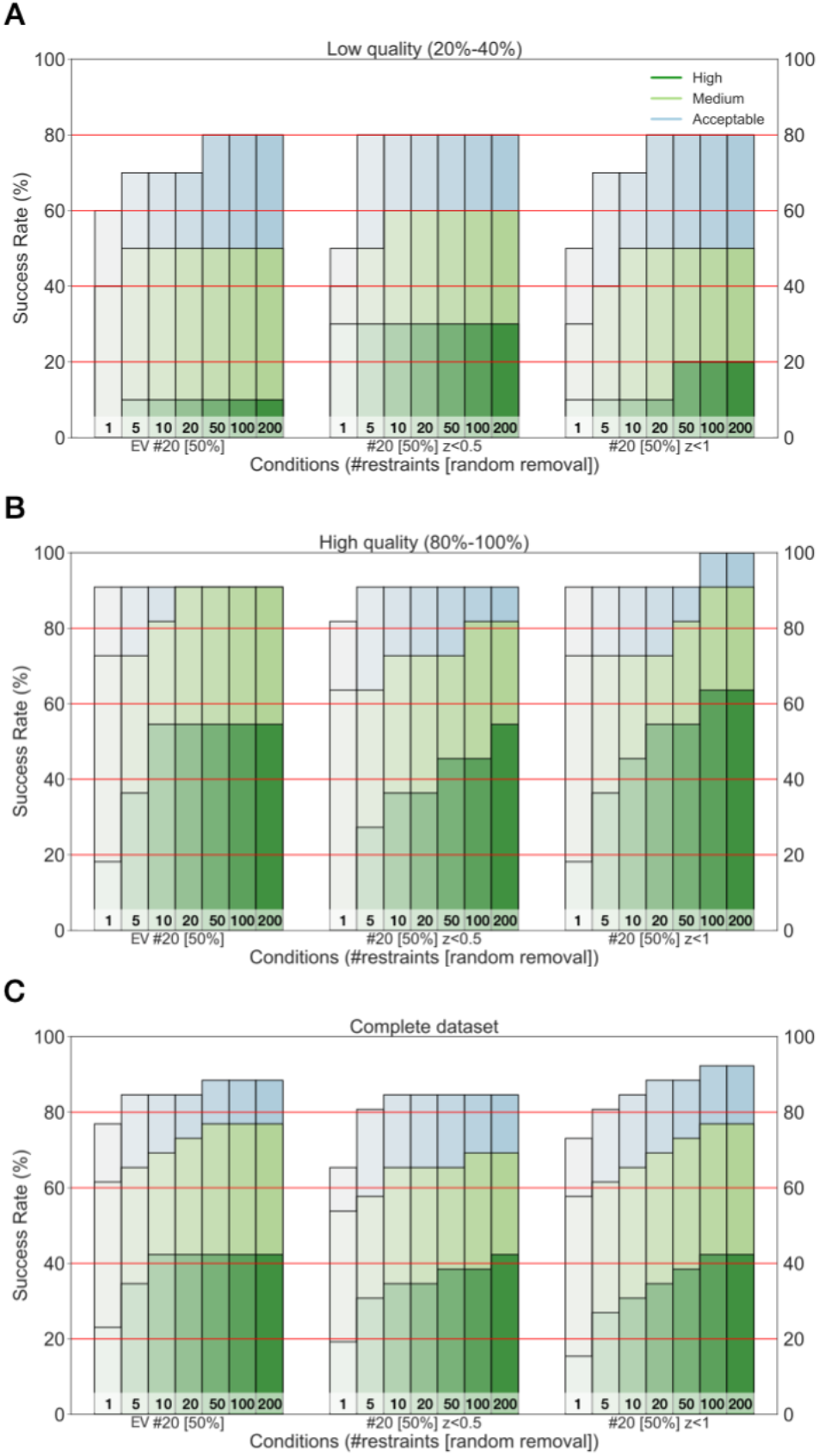
Comparison of co-evolution and DisVis-reranked docking success rates for the high- (80%-100% contact precision) and low- (20%-40% contact precision) quality datasets as well as the complete dataset of 26 complexes11 dimers of the 26 dimers dataset, including complexes with 8, 9 and 10 contacts in their Green top 10 (Table 1). **(A)** Low-quality sSuccess rate of co-evolution (EV #20[50%]) and DisVis-reranked (#20[50%] z<0.5 and #20[50%] z<1) contact lists used as input for protein-protein docking. Three sets of contact lists, 5, 10 and 20, were used to assign distance restraints in HADDOCK. Seven bars have been plotted per condition, denoting the top 1, 5, 10, 20, 50, 100 and 200 structures according to the HADDOCK itw score. The assignment of a high, medium or acceptable label to a protein complex represents its accuracy in iRMSD with high being ≤ 1 Å (dark green), medium ≤ 2 Å (light green) and acceptable ≤ 4 Å (light blue). **(B)** High-quality sSuccess rate of co-evolution (EV #20[50%]) and DisVis-reranked DisVis contact lists (#20[50%] z<0.5 and #20[50%] z<1) used as input for protein-protein docking. Three sets of contact lists, 5, 10 and 20, were used to assign distance restraints in HADDOCK. Seven bars have been plotted per condition, denoting the top1, 5, 10, 20, 50, 100 and 200 structures according to the HADDOCK itw score. The assignment of a high, medium or acceptable label to a protein complex represents its accuracy in iRMSD with high being ≤ 1 Å (dark green), medium ≤ 2 Å (light green) and acceptable ≤ 4 Å (light blue). **(C)** Success rate of co-evolution (EV #20[50%]) and DisVis-reranked DisVis contact lists (#20[50%] z<0.5 and #20[50%] z<1) used as input for protein-protein docking including the complete dataset of 26 complexes. Seven bars have been plotted per condition, denoting the top1, 5, 10, 20, 50, 100 and 200 structures according to the HADDOCK itw score. The assignment of a high, medium or acceptable label to a protein complex represents its accuracy in iRMSD with high being ≤ 1 Å (dark green), medium ≤ 2 Å (light green) and acceptable ≤ 4 Å (light blue).

**Supporting Table 2.** Average number of contacts and contact precision in complete dataset as well as low- and high-quality subsets. The three conditions that are described are the original top20 (EV20), the DisVis-reranked top20 with z-scores higher than 0.5 removed (DisVis20<0.5) and the DisVis-reranked top20 with z-scores higher than 1.0 removed (DisVis20<1.0).

| Protocol | Average #contacts | Average precision (%) |
| --- | --- | --- |
| **Complete dataset (n=26)** |  |  |
| EV20 | 19.2 ±1.2 | 48 ±25 |
| DisVis20<0.5 | 10.6 ±2.7 | 68 ±29 |
| DisVis20<1 | 16.6 ±2.7 | 55 ±32 |
| **Low-quality subset (n=10)** |  |  |
| EV20 | 18.3 ±1.5 | 27 ±11 |
| DisVis20<0.5 | 8.8 ±1.0 | 51 ±27 |
| DisVis20<1 | 17.7 ±2.5 | 26 ±12 |
| **High-quality subset (n=11)** |  |  |
| EV20 | 19.8 ±0.4 | 65 ±23 |
| DisVis20<0.5 | 12.6 ±2.9 | 83 ±30 |
| DisVis20<1 | 15.9 ±2.8 | 79 ±28 |

**Supporting Figure 6.**
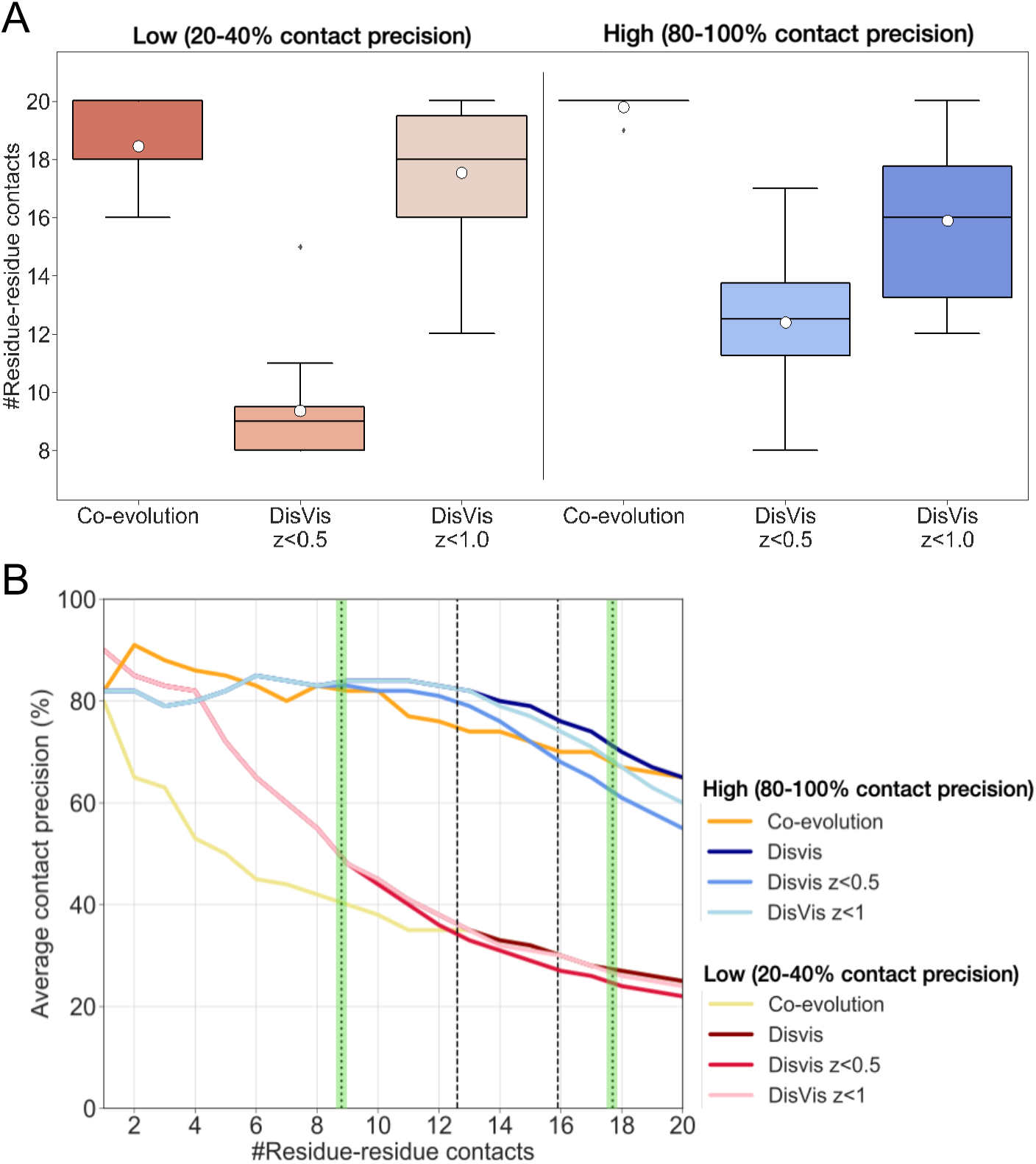
Residue-residue contact precision when considering the z-score of DisVis. **(A)** Boxplots showing total number of residue-residue contacts extracted from 20 co-evolution contacts and 20 DisVis-reranked residue-residue distances, when considering only complexes with 2 and 4 true contacts in the Green top 10 (10 dimers in low contact precision, 20-40%), or 8,9 and 10 true contacts in the Green top 10 (11 dimers in high contact precision, 80-100%), see Methods. Plots are depicted for 20-40% and 80-100% contact precision in the Green top 10, including contacts with a DisVis z-score lower than 0.5 or a z-score lower than 1.0. The mean per boxplot is depicted as a white circle. **(B)** Average contact precision for 10 dimers including 2 and 4 true contacts (20-40%) in the top 10 according to the Green top 10 (see Methods) and 80-100% true contacts in the top 10 (11 dimers). The original 20 contacts co-evolution results are shown, together with the DisVis-reranked 20 contact data (DisVis) and the DisVis results, including only contacts with a DisVis z-score lower than 0.5 or lower than 1.0. The green-highlighted dotted lines depict the mean values of the total number of contacts considered after z<0.5 removal (8.8 contacts) for the 10 dimers in 20-40% true contacts or after z<1 removal (17.7 contacts) for the 20-40% contact precision. The dashed lines represent the mean value of the total number of contacts considered after z<0.5 removal (12.6 contacts) and z<1 removal (15.9 contacts) for the 11 dimers with 80-100% true contacts (Supporting Table 2).

